# PAK4 regulates stemness and progression in endocrine resistant ER-positive metastatic breast cancer

**DOI:** 10.1101/558528

**Authors:** Angélica Santiago-Gómez, Ilaria Dragoni, Roisin NicAmhlaoibh, Elisabeth Trivier, Verity Sabin, Bruno M. Simões, Julia M. Gee, Andrew H. Sims, Sacha J. Howell, Robert B. Clarke

**Author notes:** Corresponding author, Breast Biology Group, Manchester Breast Centre, Division of Cancer Sciences, Oglesby Cancer Research Building, University of Manchester, Manchester, M20 4GJ, United Kingdom. CRUK Centre for Drug Development, Angel Building, 407 St John Street, EC1V 4AD, London, United Kingdom.

## Abstract

Despite the effectiveness of endocrine therapies to treat estrogen receptor-positive (ER+) breast tumours, two thirds of patients will eventually relapse due to *de novo* or acquired resistance to these agents. Cancer Stem-like Cells (CSCs), a rare cell population within the tumour, accumulate after anti-estrogen treatments and are likely to contribute to their failure. Here we studied the role of p21-activated kinase 4 (PAK4) as a promising target to overcome endocrine resistance and disease progression in ER+ breast cancers. PAK4 predicts for resistance to tamoxifen and poor prognosis in 2 independent cohorts of ER+ tumours. We observed that PAK4 strongly correlates with CSC activity in metastatic patient-derived samples irrespective of breast cancer subtype. However, PAK4-driven mammosphere-forming CSC activity increases alongside progression only in ER+ metastatic samples. PAK4 activity increases in ER+ models during acquired resistance to endocrine therapies. Targeting PAK4 with either CRT PAKi, a small molecule inhibitor of PAK4, or with specific siRNAs abrogates CSC activity/self-renewal in clinical samples and endocrine-resistant cells. Together, our findings establish that PAK4 regulates stemness during disease progression and that its inhibition reverses endocrine resistance in ER+ breast cancers.

**Highlights:**

- PAK4 predicts for failure of endocrine therapies and poor prognosis
- PAK4 drives stemness and progression in ER+ metastatic breast cancer
- Targeting PAK4 abrogates breast CSC activity and restores sensitivity to endocrine treatments
- Targeting PAK4 will improve outcome of ER+ breast cancer patients

**List of Abbreviations that appeared in abstract:** Cancer Stem-like Cells (CSCs)

p21-activated kinase 4 (PAK4)

Estrogen Receptor (ER)

## 1. Introduction

Endocrine resistance is a major problem for the treatment of Estrogen Receptor (ER)-positive breast tumours. Despite their undoubted benefit in clinical practice, anti-estrogen therapies fail for at least two thirds of ER+ breast cancer patients due to *de novo* or acquired resistance, which eventually lead to metastatic relapse [1]. Several studies have reported that Cancer Stem-like Cells (CSCs) are enriched following endocrine therapies [2-4]. This rare population of cancer cells with stem-like features and tumour-initiating ability is enriched by radio-, chemo- and endocrine therapies, and likely to be responsible for their failure and subsequent disease progression [4-6]. Different molecular mechanisms account for the development of endocrine resistance, which mainly revolve around ER function. In fact, ER expression is absent or low in breast CSCs [7]. In addition to the loss of ER, other mechanisms are the acquisition of gain-of-function mutations in *ESR1* [8-11] or expression of truncated ER variants [12] as disease progresses to an advanced state. Moreover, aberrant expression of cell cycle regulators that counteract the cytostatic effect of anti-estrogens or the deregulation of receptor tyrosine kinase signalling (e.g. overexpression of epidermal growth factor family, EGFR and HER2; or insulin-like growth factor family) lead to activation of downstream pathways that can also modulate sensitivity to endocrine therapies [13-15]. These pathways have been successfully targeted by CDK4/6 and PI3K/mTOR inhibitors, leading to some benefit in ER+ patients [16-18].

p-21 activated kinases (PAKs) recently emerged as a potential druggable target to overcome endocrine resistance [19]. This conserved family of serine/threonine kinases, originally described as downstream effectors of small Rho GTPases, Rac and Cdc42, is crucial for cytoskeletal dynamics, survival, proliferation, metabolism and invasion. In mammals, six members have been identified and classified into two groups based on sequence and structure similarities: Group I, PAK1-3; and Group II, PAK4-6. PAK function is upregulated in many human cancers (including melanoma, hepatocellular carcinoma, pancreatic, ovarian, prostate and breast cancer) [20-25], and copy number aberrations have frequently been described in the chromosomal regions containing *PAK1* and *PAK4* genes [20, 21, 24, 26-28]. Data supporting a role in breast cancer include oncogenic transformation of immortalised mouse mammary epithelial cells by PAK4 overexpression and PAK4 RNAi reversing the malignant phenotype of MDAMB231 breast cancer cells [29, 30]. Moreover, 3 independent studies on the expression of PAK4 in breast clinical specimens at different disease stages showed that high protein levels correlate with larger tumour size, lymph node involvement and invasive disease [31-33]. Furthermore, PAK4 expression associates with poor clinical outcome in tamoxifen-treated patients and was demonstrated to positively regulating ER transcriptional activity in an endocrine resistant breast cancer cell line [34].

Here we show PAK4 predicts resistance to tamoxifen and poor prognosis in 2 cohorts of ER+ breast cancer tumours. Using patient-derived breast tumour cells, we demonstrate that blockade of PAK4 signalling using a small molecule inhibitor reduces CSC activity and overcomes endocrine resistance. In metastatic patients, we show PAK4 expression is associated with endocrine resistant cancer progression. Our results indicate that PAK4 is essential for maintaining CSC features in patient-derived ER+ metastatic breast cancers and in acquired resistance to endocrine therapies. We conclude that the use of anti-PAK4 therapies will help tackle resistance in ER+ breast cancer patients.

## 2. Materials and methods

### 2.1 Identification and Characterisation of CRT PAKi

Several compounds which inhibit PAK4 were identified out of a high-throughput screening on ∼80,000 small molecules from the Cancer Research UK’s Commercial Partnerships Team (formerly known as Cancer Research Technology, CRT) compound collection. Exploration of the structural-activity relationship was carried out around novel ATP competitive chemotypes, with compounds being routinely tested against both PAK4 and PAK1 (Supp. Figure 1A). “Hit compounds” were selected to progress to a cellular pharmacodynamic biomarker assay, which measured the inhibition of phosphorylation of a direct substrate of PAK4; and also, to examine toxicity by looking at drug metabolism and pharmacokinetics (DMPK) *in vitro*. Among all, CRT PAKi showed greater potency, low microsomal intrinsic clearance and reduced colony formation in a dose-dependent manner in established cell lines of different origin (Table I & II). 1 μM of CRT compound was profiled against the kinase assay panel of 456 targets (LeadHunter Panels, DiscoverX), showing a promising off-target profile. *In vivo* pharmacokinetic studies showed that its bioavailability was 49 %, and that high levels of the compound were detected in the muscle up to 7 h post-administration (Supp. Figure 1B) [23]. CRT PAKi was provided by Cancer Research UK’s Commercial Partnerships Team (London, UK).

**Table I.** *In vitro* cellular pharmacodynamics, drug metabolism and pharmacokinetics of CRT PAKi

|  |  |
| --- | --- |
| PAK4 pIC<sub>50</sub> at 20 µM ATP | 7.5 |
| PAK1 pIC<sub>50</sub> at 16 µM ATP | 8.0 |
| PD Biomarker pIC<sub>50</sub> | 6.9 |
| Microsomal Cl Int (ml/min/g liver) | 1.91 |

pIC<sub>50</sub> values in the presence of ATP, the ability of inhibiting phosphorylation of a direct substrate of PAK4 and the microsomal intrinsic clearance are shown.

**Table II.**
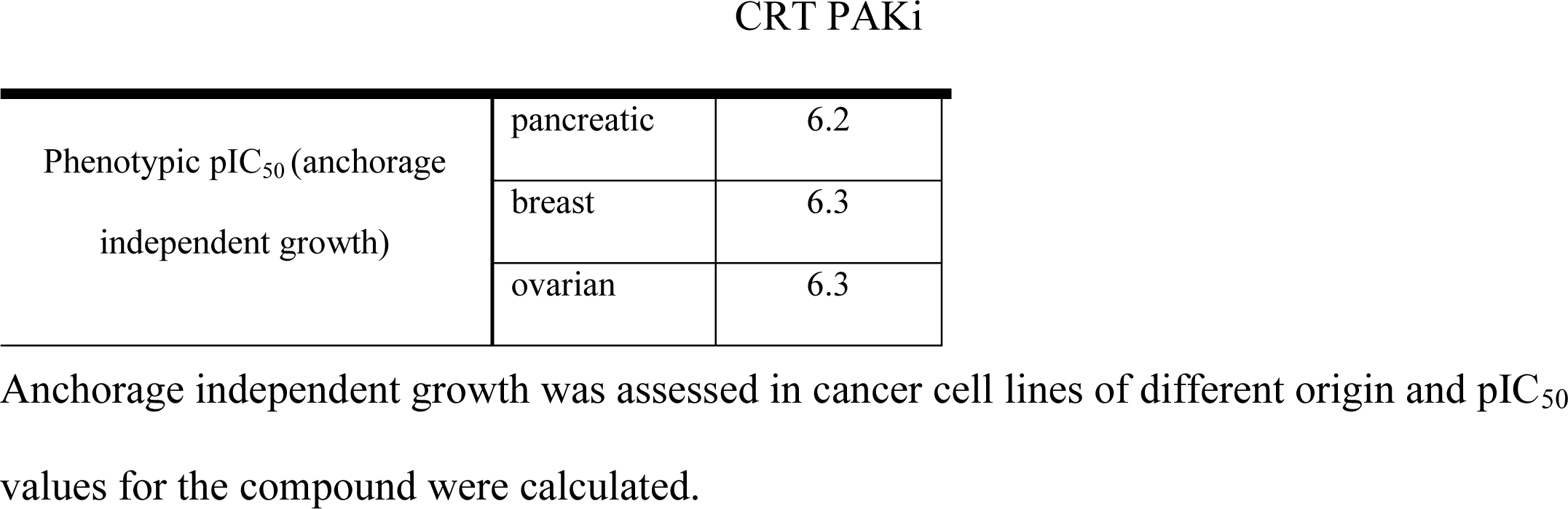
Effect of CRT PAKi on proliferation.

**Figure 1.**
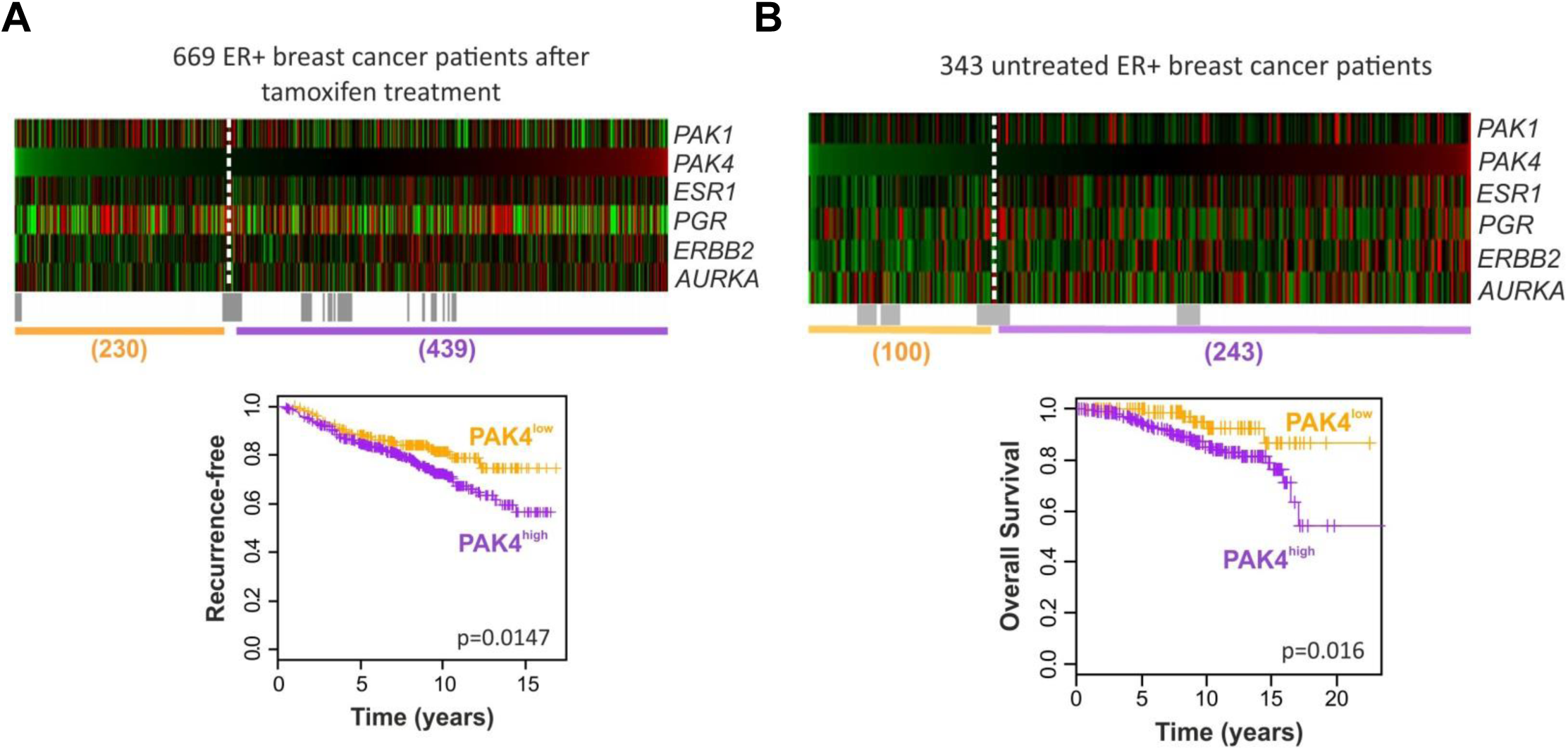
PAK4 predicts for tamoxifen resistance and poor prognosis. Expression of *PAK1, ESR1, PGR, HER2* and *AURKA* genes in 2 independent cohorts of ER+ breast cancer patients treated with tamoxifen (A) or untreated (B) is shown in the heatmaps ranked based on *PAK4* expression. Colours are log2 mean-centered values; red indicates high, whereas green indicates low expression levels. All significant cut-points (p< 0.05) are shown in grey. Kaplan-Meier analyses using the most significant cut-point (white dashed line) demonstrates that elevated expression of PAK4 is significantly associated with increased distant metastasis (A) and decreased overall survival (B).

### 2.2 Cell lines and reagents

Endocrine-resistant MCF-7 cell lines were kindly provided by Dr Julia M. Gee (University of Cardiff, Wales) [13], [35]. Parental MCF-7 cells were cultured in phenol-red DMEM/F12 media containing 10 % foetal bovine serum and 2 mM L-glutamine. Tamoxifen-resistant (TAMR) and Fulvestrant-resistant (FULVR) MCF-7 cells were routinely cultured in phenol-red DMEM/F12 media supplemented with 5% charcoal-stripped serum and 2 mM L-glutamine in the presence of either 0.1 μM 4-OH-Tamoxifen or 0.1 μM Fulvestrant, respectively. 4-OH-Tamoxifen (Sigma-Aldrich) and Fulvestrant (TOCRIS Bioscience) were purchased.

### 2.3 Metastatic patient-derived samples

Metastatic samples from breast cancer patients were collected at both The Christie NHS Foundation Trust and The University Hospital of South Manchester NHS Foundation Trust through the Manchester Cancer Research Centre Biobank (Manchester, UK). Patients were informed and consented according to local National Research Ethics Service guidelines (Ethical Approval Study No.: 05/Q1402/25 and 12/ROCL/01). Sample processing to isolate breast cancer cells from metastatic fluids (pleural effusions or ascites) was carried out as described elsewhere [36].

### 2.4 Quantitative Real-Time PCR

mRNA expression was detected using and quantitative RT-PCR. Total RNA was extracted at different conditions and qRT-PCR reactions were performed as described in [2]. Applied Biosystems Taqman Gene Expression Assays used: PAK1, #Hs000945621_m1; PAK4, #Hs00110061_m1; GAPDH, #Hs99999905_m1; and ACTB, #Hs99999903_m1. Expression levels were calculated using the ΔΔCt quantification method using *GAPDH* and *ACTB* as housekeeping genes.

### 2.5 Western blot

Cells lysates were prepared by resuspending cells in cell lysis buffer (25 mM HEPES, 50 mM NaCl, 10 % glycerol, 1 % Triton-X-100, 5 mM EDTA) containing proteases and phosphatases inhibitors (Roche MiniProtease Inhibitor cocktail; 1 μM PMSF; 30 mM sodium pyrophosphate; 50 mM sodium fluoride; 1 μM sodium orthovanadate). Then cells were incubated on rotation for 1h at 4°C, and subsequently protein lysates were obtained by centrifugation at 12,000 g at 4°C for 10 min. Protein concentration was determined using BCA Protein Assay Kit (Pierce). Samples were prepared under reducing conditions, subsequently loaded in pre-cast 10% gels (BioRad, #456-1033) and run at 200 V. Then proteins were transferred to a 0.2 μm Nitrocellulose membrane (BioRad, #170-4159) at 25 V for 15 min 1.3 A using the Transblot Turbo Transfer System (BioRad). Membranes were blocked in 5% bovine serum albumin (BSA)/ PBS-0.001% Tween 20 (PBS-T) for 1h at room temperature, followed by incubation with primary antibody diluted in 5% BSA/PBS-T at 4°C overnight. Primary antibodies used: anti-PAK1 (Cell Signaling, #2602), anti-PAK4 (Cell Signaling, #3242), β-actin (Sigma, #A2228). After 3 washes with PBS-T, HRP-conjugated secondary antibodies (Dako) were incubated for 1h at RT. Blots were developed using Luminata Classico (Millipore, Merck) and hyperfilm (Amersham GE Healthcare).

### 2.6 Mammosphere-forming assay

Cancer stem cell activity was assessed by the mammosphere-forming assay following the protocol described in [37]. When indicated, cells were directly treated in mammosphere culture with either 0.01-1 μM CRT PAKi (or control vehicle, DMSO); 1 μM 4-OH-Tamoxifen or 100 nM Fulvestrant (or control vehicle, ethanol).

### 2.7 PAK4 silencing

PAK4 expression was silenced in MCF-7 TAMR cells using siRNA technology. TAMR cells were transfected with either 10 nM control siRNA (Dharmacon, D-001810-01), siPAK4 #1 (Ambion, S20135) or siPAK4 #2 (Quiagen, SI049900000). Transfection was performed using DharmaFECT (Dharmacon) following the manufacturer’s instructions. Then transfected cells were harvested 48h post-transfection and PAK4 downregulation was confirmed by Western blot and quantitative RT-PCR.

### 2.8 Gene expression meta-analyses of ER+ primary breast tumours

The gene expression data on 669 ER+ tamoxifen-treated tumours (GSE6532, GSE9195, GSE17705, and GSE12093) and 343 ER+ untreated tumours (GSE2034 and GSE7390) was integrated from published Affymetrix microarray datasets with correction for batch effects as described previously [2]. Comprehensive survival analysis was conducted using the survivALL R package to examine Cox proportional hazards for all possible points-of-separation (low-high cut-off points) [38].

### 2.9 Statistics

Statistical significance was determined using GraphPad Prims software. Normal distribution of data was assessed using D’Agostino-Pearson, Shapiro-Wilk and Kolmogorov-Smirnov normality tests. Normal Parametric tests including one-way ANOVA with Tukey’s multiple comparisons test or two-tailed unpaired Student’s t-test were performed. When normality assumption was not possible, non-parametric Kruskal-Wallis with Dunn’s multiple comparisons test or non-parametric Mann-Whitney test were performed. Data are always expressed as mean ± SEM of at least 3 independent experiments. A p-value ≤0.05 was considered statistically significant.

## 3. Results

### 3.1 PAK4 predicts for tamoxifen resistance and poor prognosis in ER+ breast cancer

Overexpression of PAK1 and 4 in ER+ breast tumours that are refractory to endocrine therapy have previously been linked to tamoxifen resistance and poor prognosis [23, 34, 39, 40]. However, PAK4 is the only family member that associates with clinical outcome data using relapse-free survival as endpoint [34]. Then we assessed whether PAK1/4 expression would predict for patient outcome to tamoxifen treatment using overall survival data from 2 independent ER+ breast cancer patient cohorts. We carried out meta-analyses using four published Affymetrix gene expression datasets. Initially, a comprehensive survival analysis was performed on 669 pre-treated tumours of ER+ breast cancer patients, who subsequently received tamoxifen as adjuvant therapy. After ranking gene expression data by *PAK4* (low to high expression), all possible points-of-separation and their significance are shown in the survivALL plots (Supp. Figure 2A). The heatmap indicates *PAK4* expression is independent of *PAK1, ESR1, PGR, ERBB2* or the marker of proliferation *AURKA* (Figure 1A & B). At most significant cut-point, the subsequent Kaplan-Meier survival analysis revealed that high levels of *PAK4* were significantly correlated with metastatic relapse (Figure 1A, bottom panel). In contrast, only very high or very low levels of *PAK1* were associated with metastasis (Supp. Figure 2C & E). However, elevated levels of both PAKs were associated with poor clinical outcome showing reduced overall survival in an independent cohort of untreated ER+ breast cancer patients (n=343; Figure 1B, Supp. Figure 2B, D, F). Thus, PAK4 could be used as a prognostic tool to identify ER+ breast cancer patients with high risk of developing endocrine resistance and therefore benefit from the use of anti-PAK4 therapies in the adjuvant setting.

**Figure 2.**
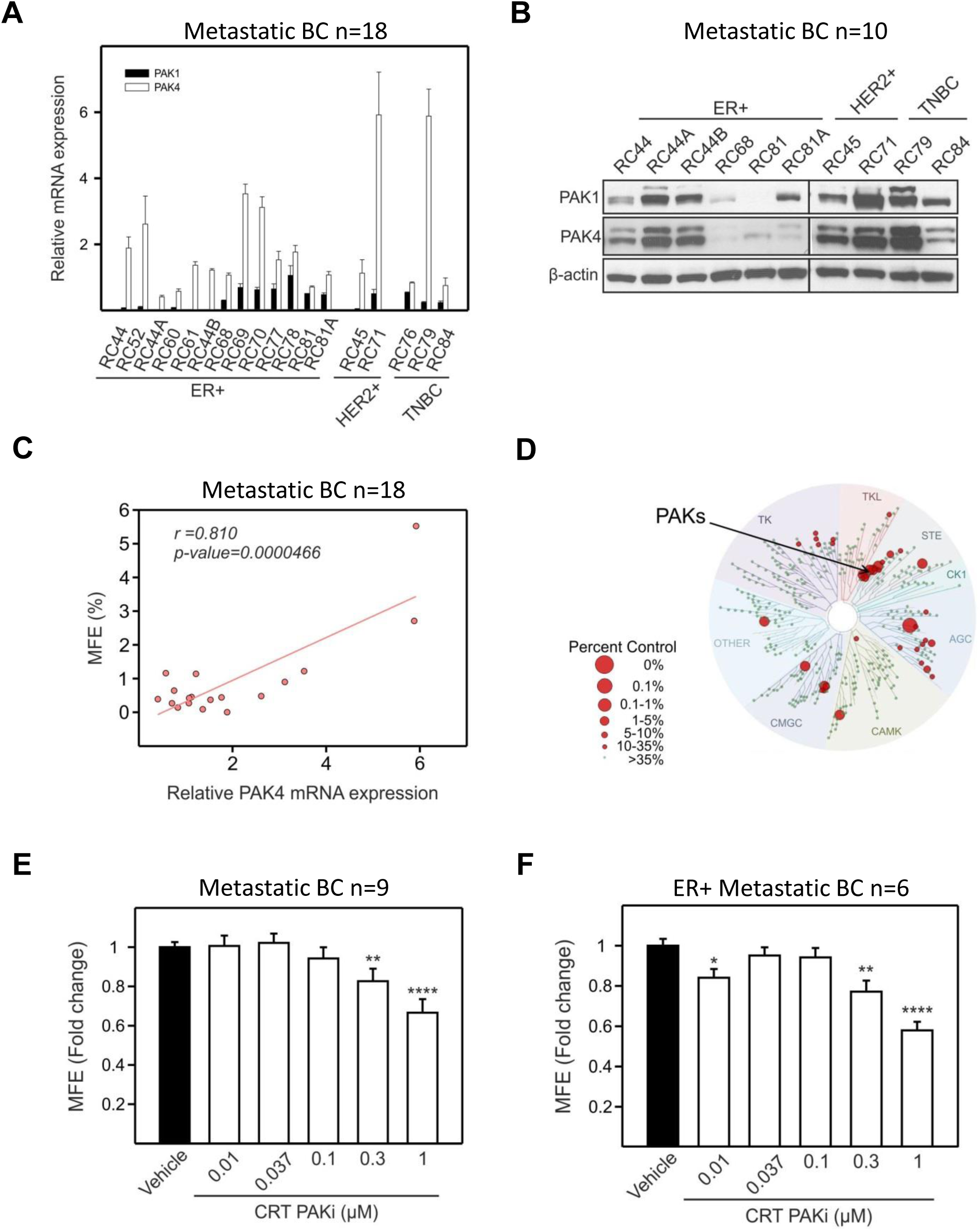
PAK4 expression correlates with stemness in metastatic breast cancer. Expression of PAK1/4 was detected at RNA (A) or protein (B) level in 18 metastatic breast cancer patient-derived samples (including all breast cancer subtypes). β-actin was used as loading control. (C) CSC activity of freshly processed metastatic samples assessed by the mammosphere-forming assay was correlated to relative PAK4 mRNA expression. (D) The off-target liability and on-target specificity of CRT PAKi is summarised in a TREE*spot*™ kinase dendrogram. 1 μM of CRT compound was screened and profiled against a kinase assay panel of 456 targets, which covers more than 80 % of human protein kinome (LeadHunter Panels, DiscoverX), using a quantitative site-directed competition binding assay [49]. In this human kinome phylogenetic tree, each kinase screened is marked with a circle. Red circles identify kinases found to bind, where larger circles show higher affinity binding; whereas small green circles indicate not significant binding. TK, non-receptor tyrosine kinases; TKL, tyrosine kinase-like kinases; STE, homologous to yeast STE7, STE11 and STE20 kinases; CK1, casein kinase 1 family; AGC, containing Protein Kinase A, G and C families; CAMK, calcium/calmodulin dependent kinases; CMGC, consists of cyclin-dependent kinases (CDK), MAPK, glycogen-synthase-3 (GSK3) and CDC-like kinases (CLK); OTHER, other kinases. Image generated using TREE*spot*™ Software Tool and reprinted with permission from KINOME*scan*®, a division of DiscoveRx Corporation, ©DISCOVERX CORPORATION 2010. (E) Overall effect of PAK1/4 inhibition using a range of concentrations of CRT PAKi on CSC activity was evaluated in metastatic patient-derived samples. (F) Detail of CRT compound effect in ER+ metastatic samples. MFE, mammosphere-forming efficiency. Stats, p-values: *≤0.05; **<0.01; ****<0.0001.

### 3.2 PAK4 expression correlates with CSC activity in metastatic breast cancer patients

PAK1/4 expression was measured in 18 patient-derived metastatic samples, including all clinically defined breast cancer subtypes (Table III, Figure 2A & 2B). We found that their expression was unrelated to subtype and that PAK4 was more frequently detected and more highly expressed that PAK1. In breast cancer cell lines, PAK4 but not PAK1 mRNA expression was significantly associated with luminal subtype (Supp. Figure 3A, B). In patient-derived samples, there was a highly significant correlation of PAK4 mRNA expression and cancer stem cell (CSC) activity measured using the mammosphere-forming assay (Figure 2C, Pearson correlation coefficient = 0.810; p-value < 0.00005; Supp. Figure 3B, Pearson correlation coefficient = 0.104; p-value = 0.682). Next, we tested the effect of increasing concentrations of a PAK1/4-specific inhibitor (CRT PAKi) on the mammosphere-forming efficiency. This compound has an encouraging off-target profile indicating high selectivity for PAK1/4 (Figure 2D). In 9 metastatic patient-derived samples PAK1/4 inhibition reduced cancer stem cell activity in a dose-dependent manner (Figure 2E). Further sub-group analysis showed this effect was due to its activity in ER+ metastatic breast cancer samples, with PAK1/4 inhibition impairing breast CSC activity up to 60 % (Figure 2F); whereas the CSC activity of triple negative samples (n=2) remained unaffected in the presence of the CRT compound (Supp. Figure 3D, E). These data suggest that PAK4 expression is important in the maintenance of the CSC pool in metastatic ER+ breast cancer.

**Table III.** Characteristics of late metastatic breast cancer patient-derived samples used in the study

| Sample | Histology | Grade | Receptor status |  |  | Chemo | Hormone Therapy | Targeted Therapy |
| --- | --- | --- | --- | --- | --- | --- | --- | --- |
|  |  |  | ER | PR | HER2 |  |  |  |
| BB3RC44 | ILC | 3 | + | + | Neg | EOX<br>Capecitabine<br>Paclitaxel | Letrozole<br>Tamoxifen<br>Exemestane<br>Fulvestrant | Herceptin<br>Lapatinib |
| BB3RC45 | IDC | 3 | + | + | + | FEC<br>Paclitaxel<br>Capecitabine<br>Vinorelbine<br>Epirubicin | Tamoxifen<br>Anastrozole | Zometa/pamidronate<br>Herceptin |
| BB3RC52 | IDC | 3 | + | + | Neg | Epirubicin<br>Vinorelbine/5FU<br>CMF<br>Docetaxel | Tamoxifen<br>Fulvestrant | Herceptin |
| BB3RC44A | ILC | 3 | + | + | Neg | EOX<br>Capecitabine<br>Paclitaxel | Letrozole<br>Tamoxifen<br>Exemestane<br>Fulvestrant | Herceptin<br>Lapatinib |
| BB3RC44B | ILC | 3 | + | + | Neg | EOX<br>Capecitabine<br>Paclitaxel | Letrozole<br>Tamoxifen<br>Exemestane<br>Fulvestrant | Herceptin<br>Lapatinib |
| BB3RC68 | IDC | 2 | + | NK | Neg | FEC-60<br>Capecitabine<br>Taxol | Tamoxifen<br>Fulvestrant |  |
| BB3RC70 | ILC | 2 | + | + | Neg | ECF | Tamoxifen<br>Letrozole<br>Anastrozole |  |
| BB3RC71 | IDC | 3 | + | + | + | FEC<br>Capecitabine<br>Vinorelbine<br>Taxotere<br>Epirubicin | Tamoxifen<br>Anastrozole<br>Fulvestrant<br>Exemestane | Herceptin<br>Lapatinib |
| BB3RC77 | ILC | ? | + | + | Neg | Paclitaxel<br>Capecitabine<br>Eribulin | Letrozole<br>Exemestane<br>Fulvestrant | Ibandronate |
| BB3RC78 | IDC | 2 | + | Neg | Neg | Epirubicin | Letrozole<br>Exemestane | Denosumab<br>Everolimus |
| BB3RC79 | IDC | 3 | Neg | Neg | Neg | FEC/Docetaxel<br>Gemcitabine/Carboplatin |  |  |
| BB3RC81 | IDC | 2 | + | + | Neg | Paclitaxel<br>Capecitabine | NK |  |
| BB3RC81A | IDC | 2 | + | + | Neg | Paclitaxel<br>Capecitabine | NK |  |
| BB3RC84 | NK | NK | Neg | Neg | Neg | AC<br>Docetaxel<br>Ixabepilone/Capecitabine<br>/Bevacizumab<br>Cisplatin/Gemcitabine | Tamoxifen |  |

<sup>1</sup> ILC, invasive lobular carcinoma; IDC, intraductal carcinoma; ER; estrogen receptor; PR, progesterone receptor; HER2, Human epidermal growth factor receptor 2; Neg, negative; EOX, Epirubicin/Oxaliplatin/Capecitabine; FEC, 5Fluorouracil/Epirubicin/Cyclophosphamide; 5FU, 5Fluorouracil; CMF, Cyclophosphamide/Methotrexate/5Fluorouracil; ECF, Epirubicin/Cisplatin/5Fluorouracil; AC, cyclophosphamide/doxorubicin; NK, not known.

**Figure 3.**
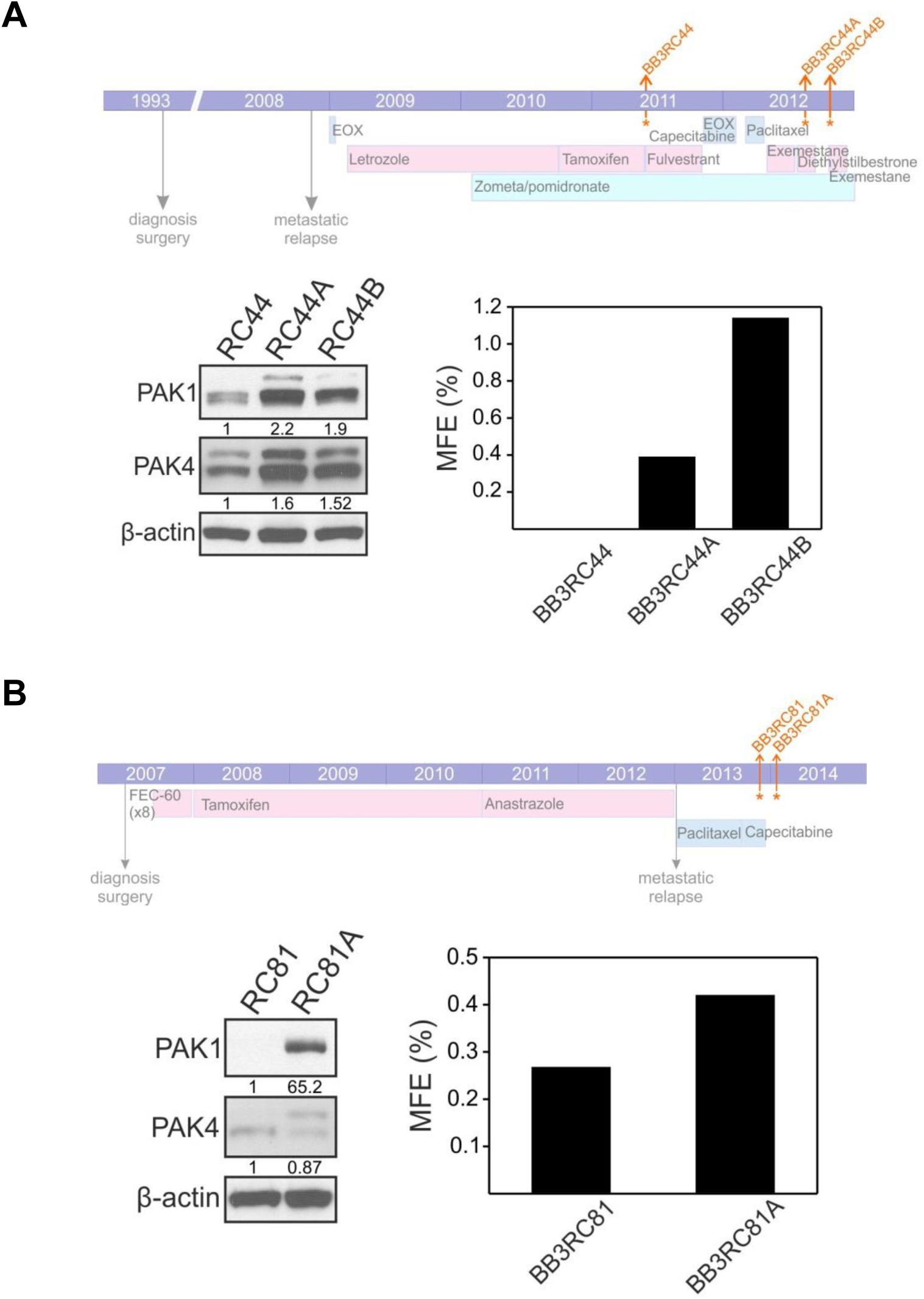
PAK1/4 and CSC activity increases during cancer progression. Examination of sequential samples of 2 ER+ metastatic patients, BB3RC44 (A) and BB3RC81 (B). The clinical treatment history of each patient is summarized in the top panels. Colours: light blue, pink or green indicate hormonal, chemo- or bone-directed therapy, respectively. Orange arrows pointed when the samples were taken. PAK1/4 protein levels and CSC activity measured as mammosphere-forming efficiency (MFE) are shown in bottom panels. Densitometric values of the ratio PAK to β-actin are indicated below the blots.

### 3.3 PAK1/4 expression is related to cancer progression

Next, we examined sequential metastatic samples of 2 ER+ breast cancer patients. The patients’ clinical treatment history is summarized in Figure 3A & B. Our analyses showed that both PAK1/4 protein levels and CSC activity increased alongside with disease progression. We detected increased expression of both PAK family members in samples from patient BB3RC44 (∼2 or 1.6-fold for PAK1/4, respectively, Figure 2A), whereas a striking increase of PAK1 levels was observed in patient BB3RC81 (∼65-fold, Figure 2B). These results show that an increase in PAK1/4 expression is correlated with disease progression in ER+ breast cancers, establishing their involvement in the failure of endocrine therapies.

### 3.4 PAK4 downregulation restores endocrine sensitivity in resistant cells

These patient data suggest either PAK1 or −4 or both have a role in endocrine resistance. To test our hypothesis, we used *in vitro* ER+ MCF-7 cell lines of acquired resistance after long-term exposure to either tamoxifen (TAMR) or fulvestrant (FULVR), respectively [13, 35]. Initially, we assessed the expression of PAK1/4 in parental, TAMR and FULVR cells. Both PAK1 and PAK4 were overexpressed in resistant cells compared to parental cells (Figure 4A). However, short-term treatment with tamoxifen or fulvestrant in MCF-7 cells did not induce a significant upregulation of *PAK1/4* gene expression (data not shown). To further confirm the role of PAK1/4 in endocrine resistance and stemness, we evaluated CSC activity for endocrine resistant cells in the presence of CRT PAKi. PAK1/4 chemical inhibition abrogated CSC self-renewal (>95 % in TAMR and 80 % in FULVR, respectively; Figure 4B) but not primary mammosphere formation (Supp. Figure 4A). Similarly, PAK4 silencing in TAMR cells not only impaired breast CSC activity (Figure 4C & Supp. Figure 4B), but also restored their sensitivity to tamoxifen (Figure 4D). These findings indicate that breast CSC activity in endocrine resistant cells depends on PAK4, which can be targeted to overcome endocrine resistance. We hypothesized that PAK4 inhibition in combination with endocrine therapies will benefit ER+ breast cancer patients. To test this, we treated 4 ER+ patient-derived breast cancer metastatic samples with either fulvestrant or CRT PAKi as single agents or in combination. CRT PAKi on its own did not have a significant effect on MFE but its combination with the standard of care fulvestrant had a synergistic effect reducing CSC activity more than half. When patient-derived samples were separated into responders versus non-responders, we identified that only ER+ breast cancer patients with high levels of PAK4 benefit from the combination of therapies (Figure 4E and Supp. Figure 4C), suggesting PAK4 expression is a predictive biomarker of response. These results confirm the importance of targeting PAK4 to potentiate endocrine therapy and overcome resistance.

**Figure 4.**
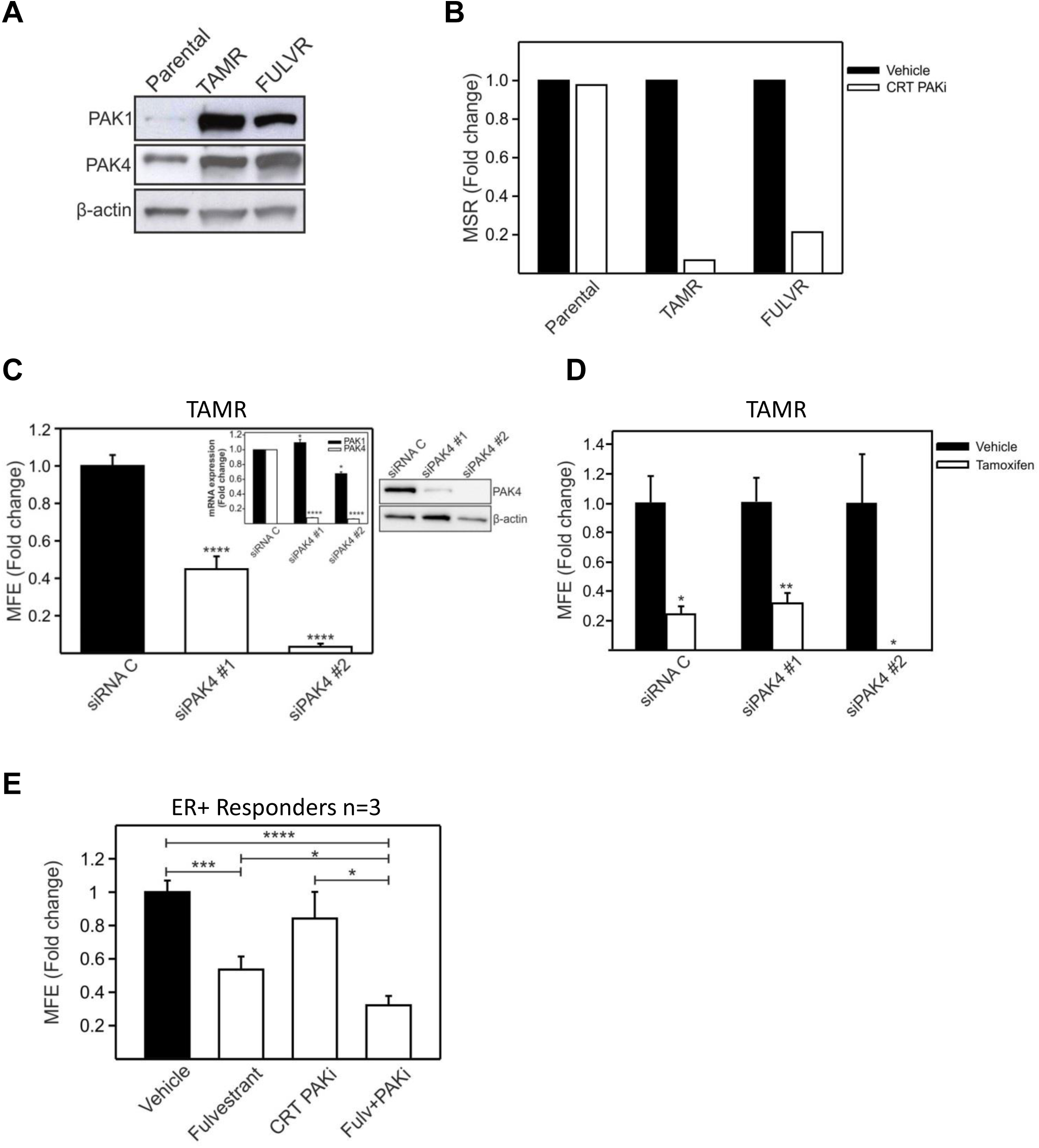
PAK4 downregulation restores anti-estrogen sensitivity in resistant cells. (A) PAK1/4 expression was detected in endocrine resistant MCF-7 cells by Western Blot. (B) Second generation mammospheres were plated to assess for mammosphere self-renewal (MSR) of cells treated in the first generation with 0.5 μM CRT PAKi or vehicle (DMSO) in resistant cells. (C) Effects of PAK4 silencing on CSC activity in TAMR cells. Two different siRNAs were used against PAK4 (siPAK#1, siPAK#2). Then CSC activity of siRNA-transfected TAMR cells was evaluated using the mammosphere-forming assay. The inset shows PAK1/4 mRNA expression in siRNA-transfected cells. In the right panels, PAK4 downregulation at protein level was observed by Western Blot. (D) PAK4-silenced TAMR cells were cultured with either 1 μM tamoxifen or vehicle control (ethanol) during the mammosphere-forming assay. Mammosphere-forming efficiency (MFE) is shown. (E) Combination of PAK4 inhibition and anti-estrogen therapies in ER+ metastatic breast cancer. Mammosphere-forming efficiency of patient-derived samples treated with either 0.5 μM CRT PAKi, 100 nM fulvestrant or both therapies was assessed. β-actin was used as loading control. Stats, p-values: * ≤0.05; **<0.01; ***<0.001; ****<0.0001.

## 4. Discussion

Despite the remarkable impact on survival caused by the introduction of endocrine therapies for the treatment of ER+ breast cancers, late recurrences occur in some patients due to the development of resistance to these single agents. Several authors have shown that breast CSC activity and frequency are enhanced upon endocrine therapies such as tamoxifen and fulvestrant, suggesting that this drug-resistant population accounts for the eventual metastatic relapse [2, 3]. Here we report for first time that PAK4 signalling is essential for maintaining CSC features in ER+ metastatic breast cancers. Also, PAK4 can be used as a predictive biomarker of response to endocrine therapies, and furthermore, its inhibition reverses endocrine-driven resistance in ER+ breast cancer patients.

The relationship between PAK4 and stemness has previously been described in pancreatic cancer cell lines [41, 42]. In this study, pancreatic CSCs express high levels of PAK4 and its silencing reduced not only sphere formation, but also stem cell-related markers [42]. In agreement with these findings, we found that PAK4 significantly correlated with mammosphere-forming ability, and treatment with CRT PAKi reduced breast CSC activity in a dose-dependent manner in metastatic samples of all subtypes. Using RNA-seq data from 10 breast cancer Patient-Derived Xenografts (PDXs), we observed that *PAK4* expression correlated with *DLL1, NOTCH1-4, PTCH* and *GLI1* (data not shown). These genes are involved in NOTCH or Hedgehog signalling, both developmental pathways that regulate CSC homeostasis and self-renewal [43].

Most importantly, the effect of PAK4 inhibition on CSCs is restricted to ER+ metastatic samples, as the presence of CRT PAKi did not alter CSC activity of ER-subtype. In fact, *PAK4* expression significantly correlated with stem cell-related genes such as *SOX2, POU5F1* or *ALDH1A3* only in metastatic ER+ PDXs (data not shown). PAK4 is often amplified in basal-like cancers, which give rise to TNBC [26]; and silencing PAK4 or using inhibitors that induce protein destabilisation reduce proliferation and *in vivo* tumorigenesis in TNBC, but not in ER+ or HER2+ cell lines [29, 44]. This discrepancy in the role of PAK4 between breast cancer subtypes might be either associated with its additional kinase-independent functions [33, 45], which are compromised upon reducing protein levels and therefore could drive tumorigenesis in TNBC; or, instead, with off-target activity of these inhibitors, e.g. affecting enzymes involved in NAD metabolism [46]. Mechanistically, differences among subtypes can be related to the presence of ER, as a positive feedback loop has been described where ER promotes PAK4 expression and, in turn, PAK4 regulates its transcriptional activity in endocrine resistant cells [34]. Further investigation is needed to fully understand the specific resistance mechanism in each breast cancer subtype. In most adult tissues, PAK4 expression is low. However, its overexpression has not only been associated with oncogenic transformation [29, 30], but also with disease stage in breast clinical specimens [31-33]. We found that PAK1/4-driven CSC activity increased as the disease progressed in sequential metastatic samples taken from 2 ER+ breast cancer patients. However, PAK4 expression only increased during progression in patient BB3RC44, who received several lines of endocrine therapy after metastatic relapse, suggesting a resistant phenotype. Whereas patient BB3RC81 was treated with just chemotherapy after recurrence and progression seems to rely on a PAK1-dependent mechanism.

Then we confirmed overexpression of PAK4 in endocrine resistant MCF7 cells. Importantly, CRT PAKi abrogated almost completely CSC self-renewal and silencing of PAK4 not only reduced mammosphere formation, but also restored the effect of tamoxifen in endocrine resistant cells. Restoration of sensitivity has already been reported using GNE-2861, a group II PAK inhibitor, in tamoxifen-resistant MCF7/LCC2 cells [34]. Furthermore, blocking PAK4 in combination with standard of care fulvestrant reduced CSC activity even further in ER+ breast cancer metastatic samples with high levels of PAK4. Therefore, PAK4 not only has prognostic value as confirmed using overall survival data as clinical end point; but it is also a predictive biomarker of response to endocrine therapies. Thus ER+ breast cancer patients with high levels of PAK4 could be identified and benefit for using PAK-targeting therapeutics. In recent years, considerable efforts have been made to develop PAK inhibitors. PF-3758309, which targets group I and II PAKs, was the first PAK inhibitor to enters clinical trials for advanced solid tumours. Although it blocks growth of a variety of tumour cell lines *in vitro* and *in vivo*, it failed in phase I due to adverse pharmacological properties and side effects [47]. Since then, many attempts have been made to develop novel small molecules inhibitors with good oral bioavailability [48]. KPT-9274 is currently in phase I clinical trials for solid tumours and lymphomas. The inhibitory mechanism and off-target effects of this destabilising agent still remain to be elucidated. Although it has been reported to be promising for controlling tumour growth in TNBC, ER+ and HER2+ cell lines are unresponsive [44]. Therefore, other strategies must be considered to target ER+ disease. Here we showed that blocking only kinase-dependent functions of PAK4 using CRT PAKi is sufficient to overcome endocrine resistance.

In conclusion, we report for first time that PAK4 is a promising target to reduce CSC activity in ER+ metastatic breast cancers and furthermore its expression can be used as a prognostic and preventive tool for patient stratification to identify those who will benefit from complementary anti-PAK4 therapies.

## 6. Acknowledgements

A.S.G. was recipient of a Postdoctoral Fellowship granted by the Alfonso Martín Escudero Foundation. We are grateful for funding from Cancer Research UK and Breast Cancer Now (Grants Nos. 2015NovPR651 and MAN-Q2-Y4/5).

## 8. Supplementary material /Additional information

Supplementary file

Supplementary Figure legends

