## Supplemental Figure Legends for "PAK4 regulates stemness and progression in endocrine resistant ER-positive metastatic breast cancer"

### Supplementary Figure Legends

**Supplementary Figure 1. Identification and characterization of CRT PAKi.** (A) Two series of compounds (yellow and orange) were tested against both PAK1 and PAK4. $pIC_{50}$ values for each kinase are represented. (B) Pharmacokinetic profile of CRT PAKi was tested in female CD-1 mice. 30 mg/kg of the compound were orally administered in 1 dose and then its concentration was measured in muscle and blood overtime.

**Supplementary Figure 2. Prognostic value of PAKs.** All possible points-of-separation across 2 independent cohorts of ER+ breast cancer patients (tamoxifen-treated, n= 669; untreated, n= 343) are shown in the survivALL plots (A-D). Patients were ranked from low to high expression of *PAK4* (A-B) or *PAK1* (C-D) and all hazard ratio for all possible cut-points were displayed. Colours indicate the significance of each point-of-separation (low-high significance= blue-yellow gradient). (C-F) Prognostic value of *PAK1*. The heatmaps show the expression of *PAK4*, *ESR1*, *PGR*, *ERBB2* and *AURKA* genes ranked by *PAK1* expression in the tamoxifen-treated (E) or untreated (F) cohorts. Colours are log2 mean-centred values; red indicates high, whereas green indicates low expression levels. All significant cut-points ($p < 0.05$) are shown in grey. Kaplan-Meier analyses using the most-significant cut-point (white dashed line) demonstrates that either very high or very low expression of PAK1 is significantly associated with increased distant metastasis (E), whereas only high expression associates with decreased overall survival (F).

**Supplementary Figure 3. PAK expression in breast cancer.** Box plots show the gene expression of *PAK4* (A) or *PAK1* (B) across datasets of 51 breast cancer lines available through the GOBO online tool [38]. In the left panel, data was divided in Basal A (blue),

Basal B (grey) and Luminal (red); whereas in the right panel, data was grouped into clinical subtypes: TN, triple negative (blue); HER2, HER2-positive (purple); and HR, Hormone receptor positive (red). Classification based on [39]. (C) CSC activity of freshly processed metastatic samples assessed by the mammosphere-forming assay and correlated to relative PAK1 mRNA expression. Inhibition of PAK1/4 by a range of concentrations of CRT PAKi on CSC activity was determined in HER2+ (D) or TN (E) metastatic patient-derived samples. Mammosphere-forming efficiency (MFE).

Stats, p-values: *≤0.05; ****<0.0001.

**Supplementary Figure 4. Effect of CRT PAKi and PAK4 silencing on CSC activity in endocrine resistance.** (A) CSC activity of endocrine resistant MCF7 cells (parental, TAMR and FULVR) was evaluated in the presence of 0.5 μM CRT PAKi or vehicle control (DMSO) by the mammosphere-forming assay. Mammosphere-forming efficiency (MFE) normalized to vehicle is shown. (B) PAK1 levels were detected by Western Blot in siPAK4-transfected TAMR cells. (C) Combination of PAK4 inhibition and anti-estrogen therapies in a non-responder ER+ metastatic breast cancer. Mammosphere-forming efficiency of the patient-derived sample cultured with either 0.5 μM CRT PAKi, 100 nM fulvestrant or both therapies was assessed.
