## Supplemental Figures for "PAK4 regulates stemness and progression in endocrine resistant ER-positive metastatic breast cancer"

### Supplementary info – Figure 1

A

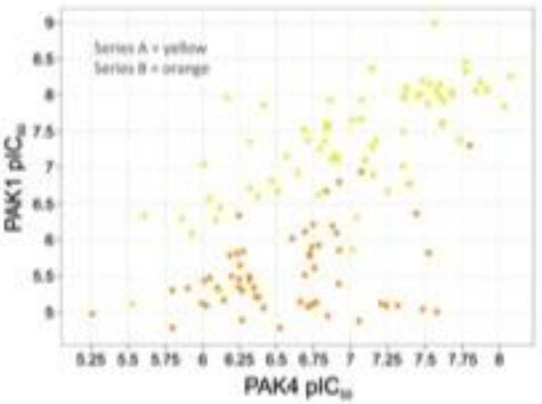


B

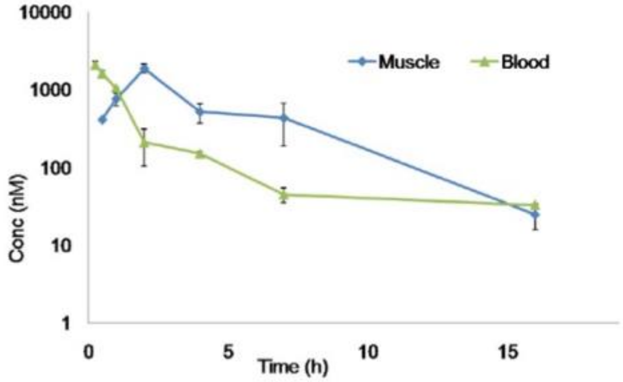

### Supplementary info – Figure 2

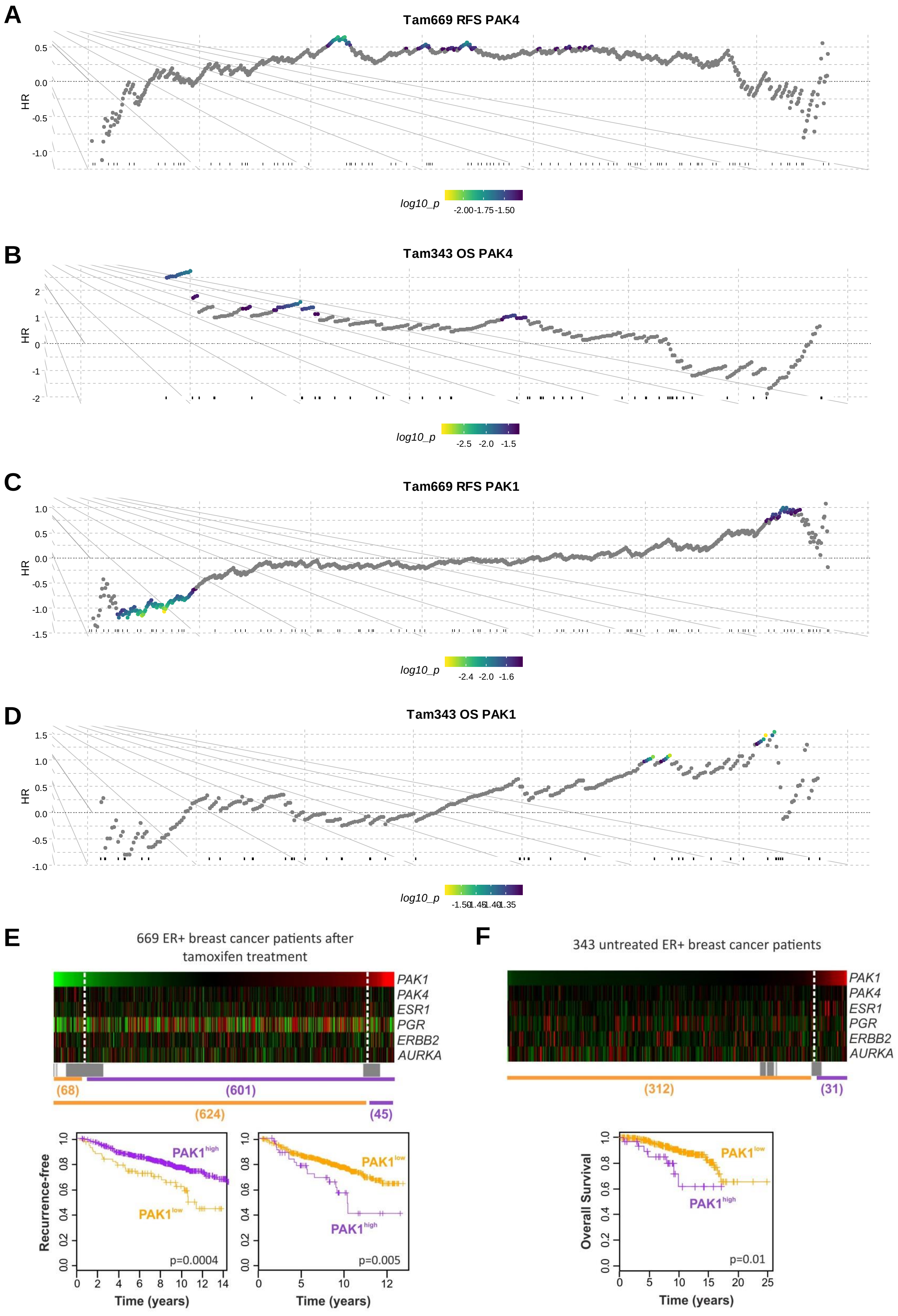

### Supplementary info – Figure 3

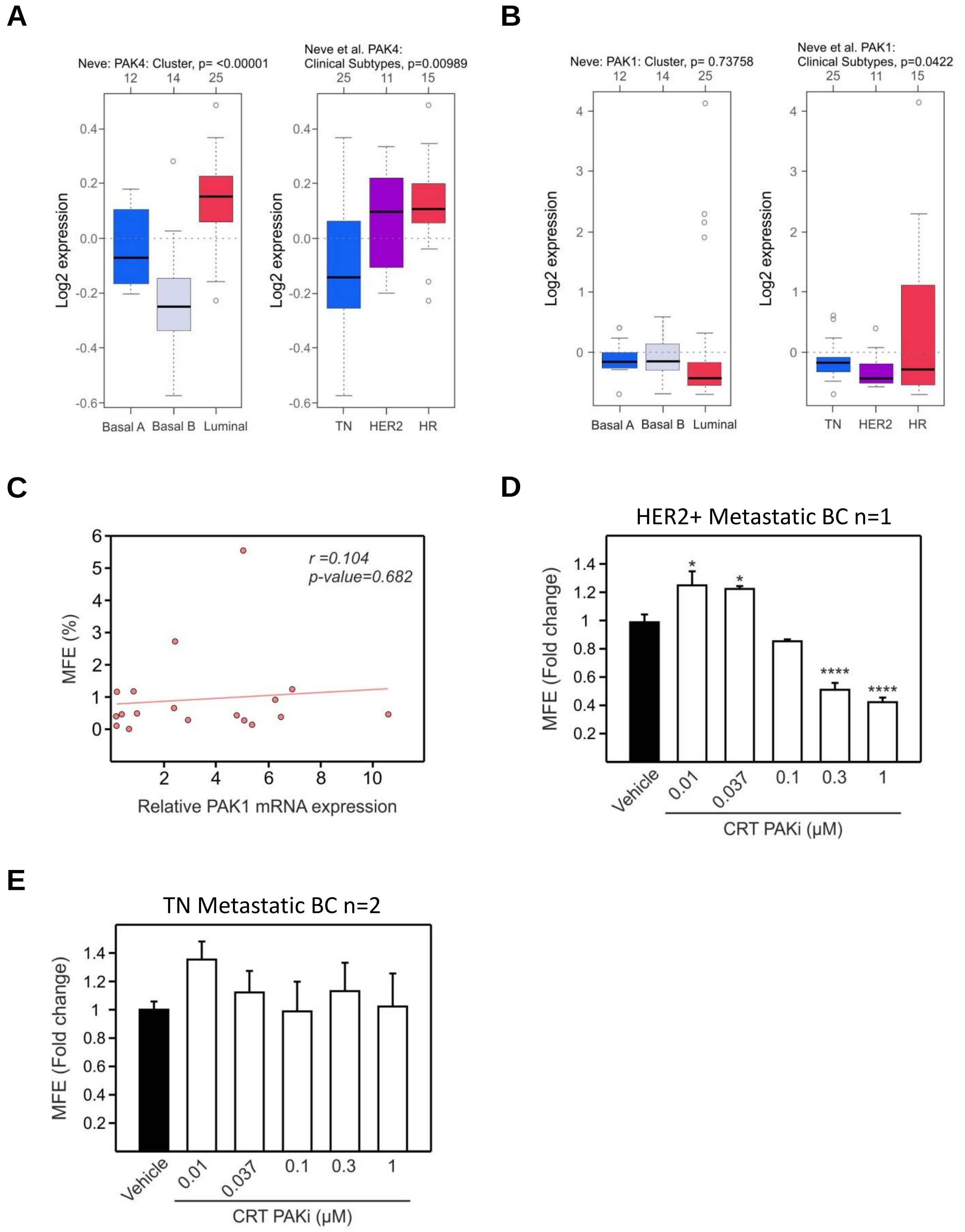

### Supplementary info – Figure 4

A

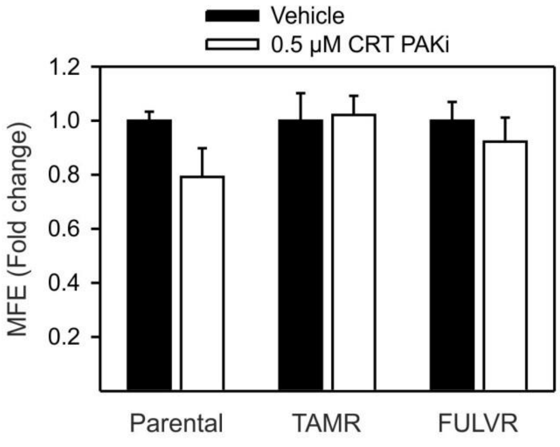


B

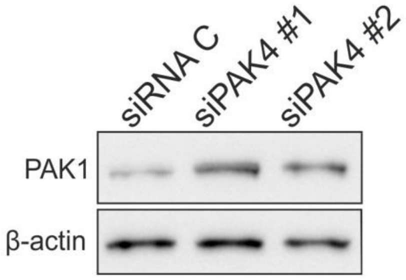


C

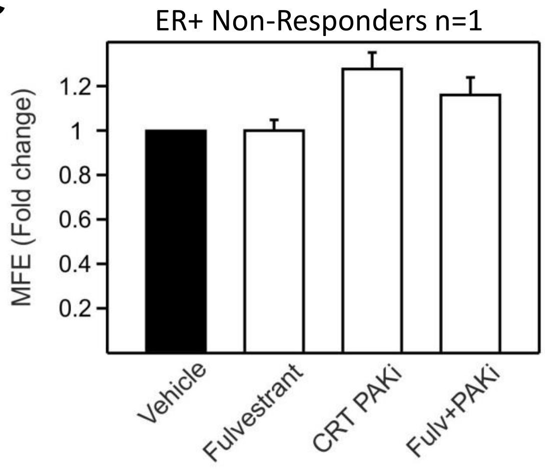
